# Expression and molecular regulation of key genes during adipogenesis of neural crest stem cells

**DOI:** 10.1101/2023.06.07.544147

**Authors:** Bo-wen Dong, Wen-chao Guan, Kai Zhang, Yan Zhang, Yu Yang, Yan-ping Zhao, Rui Bai, Ming-xue Zheng, Xiao-zhen Cui

## Abstract

Neural crest stem cells (NCSCs) are pluripotent stem cells derived from the “fourth germ layer”. Similar to mesenchymal stem cells (MSCs), NCSCs can differentiate into a variety of cell types, such as adipocytes. However, the mechanism of NCSCs adipogenesis remains unclear. Previously, we have revealed that primary cells have neural crest and stem cell properties and can differentiate into adipocytes. Therefore, in this study, the differentially expressed genes (DEGs) of NCSCs at specific time points of adipogenesis were predicted by mRNA sequencing, the key genes of adipogenesis were predicted by principal component analysis, heat map, GO and KEGG enrichment analysis, and the expression of DEGs was verified by RT-qPCR. RNA interference (RNAi) technology was used to inhibit the expression of DEGs, and RT-qPCR and Western blot were used to explore the regulatory mechanism between DEGs. Sequencing results indicated a possible regulatory relationship between C/EBPα, C/EBPβ, C/EBPδ and PPARγ. The results of RT-qPCR were consistent with those of mRNA sequencing. Combined with RT-qPCR and western blot results, we found that C/EBPβ and PPARγ regulated the transcription of C/EBPα during NCSCs adipogenesis, and C/EBPβ and PPARγ formed positive feedback loop.

## 1. INTRODUCTION

Neural crest cells (NCCs) are a group of temporary cells separated from the neural crest (NC) called the fourth germ layer [1], which is a special structure in the early embryonic development of vertebrate. NCCs are strong plasticity in development and can produce a variety of “nerve” derivatives including melanocytes, craniofacial cartilage and bone, smooth muscle, peripheral and enteral neurons and glial cells [2,3,4]. As their pluripotency, Stemple et al named them neural crest stem cells (NCSCs) [5]. Migration of NCCs is a highly coordinated process. After the neural folds close to form the neural tube, the cells located at the edge of the neural plate transform into NCCs and begin to migrate along the outer side of the neural tube. After epithelial-mesenchymal transition (EMT) [6], NCCs will migrate and differentiate into various tissues and organs. Thus, NCCs has broad application prospects in tissue engineering, but the difficulty of obtaining large quantities of NCCs is the main factor limiting its application in regenerative medicine research.

However, the emergence of adipose-derived stem cells have broken this impasse. The ease of extracting adipose tissue without ethical constraints makes it an attractive candidate tissue source for medical applications [7]. NCSCs have been isolated from human subcutaneous adipose have been reported. In addition, NCCs can be differentiated into brown and white adipose cells in vitro and in vivo [8]. Moreover, NCCs derived adipose tissue and non-NCCs derived adipose tissue have different transcriptional profiles [9]. Another study isolated and identified a population of adipose-derived stem cells with NCSCs characteristics from adult bovine adipose tissue and named them as bovine adipose-derived neural crest stem cells (baNCSCs) [10]. Therefore, NCSCs derived from subcutaneous adipose have great research potential and are rare candidate cells for medical application.

The 3T3-L1 cell line as model of differentiation of adipocytes have been widely reported [11]. Mitotic clonal expansion is required step in the adipocyte differentiation program, but is not necessary for 3T3-L1 preadipocytes and C3H10T1/2 cells [12,13]. At this point, cells begin to transform from long spindle into irregular polygonal shape and increase in size [14]. With the completion of mitosis, the preadipocytes exit the cell cycles and enter terminal differentiation. In the processes, Krox20 induces CCAAT/enhancer binding protein β (C/EBPβ) expression [15] and binds to cAMP-response element binding protein (CREB) regulates C/EBPβ expression [16]. C/EBPβ and C/EBPδ activate Kruppel like factor 5 (KLF5), and then collectively induce peroxisome proliferator-activated receptor γ (PPARγ) [17], PPARγ activates C/EBPα with C/EBPβ and C/EBPδ, constituting the classical C/EBPβ+δ-PPAR-C/EBPα pathway [18].

Adipose differentiation and development are important trait widely studied at home and abroad, which not only has economic value, but also has important medical value for the study of obesity and other diseases. Pigs and humans are similar regulatory patterns during adipose differentiation and development. However, the regulatory mechanism of porcine subcutaneous adipose NCSCs differentiation into adipose lineage has not been reported. In this research, the primary cells were identified by neural crest and stem cell markers. The key factors and pathways of adipogenesis were predicted by mRNA sequencing. The regulatory relationship between key adipogenic factors related to this pathway and the adipogenic differentiation of porcine subcutaneous adipose NCSCs was explored. It provides a new direction for the clinical treatment of fat deposition, diabetes, fatty liver and other fat accumulation diseases.

## 2. MATERIALS AND METHODS

### 2.1 Ethics Statement

The animals used in all experiments were reviewed by the Experimental Animal Ethics Committee of Shanxi Agricultural University, and the experimental animal use protocol was in accordance with the ethical requirements of animal experiments.

### 2.2 Isolation and maintenance of paNCSCs

Pig subcutaneous adipose tissue was collected from the randomly selected healthy piglets (5 males and 5 females), the tissues were washed with High Glucose DMEM (Hyclone) supplemented with penicillin/streptomycin (50 units/mL, Gibco), and dissected to small pieces (less than 0.2 cm diameter). The pieces of adipose tissue were explanted in collagen-coated 3.5 cm diameter culture dishes added with the above described medium additionally supplemented with fetal bovine serum (10%, Gibco), sodium pyruvate (1mM, Sigma), uridine (50 mg/mL, Sigma), and amphotericin B (1.25 ug/mL, Gibco) as growth medium, which enables medium to inmmerse the tissue pieces. A sterilized cover slip was used to cover the tissue, cultured at 37°C in an incubator with 5% CO2/ 95% air, changing the medium every 4 days. Migratory cells with 70-80% confluence were separated with accutase (Sigma), and passaged them into a new culture dishes. The culture conditions were the same as the primary culture

### 2.3 Immunofluorescence

The cells were washed with PBS twice for 5 minutes each time. Cells were fixed with 4% paraformaldehyde for 15 minutes at room temperature, subsequently permeabilized with 0.25% Triton X-100 for 15 minutes. Following three times washing, cells were blocked with 10% goat serum and 10% donkey serum in PBS for 30 minutes respectively, and incubated with p75NTR primary antibodies (1:200, Bioss) and SOX2 primary antibodies (1:200, Abcam) respectively overnight at 4°C. The Goat Anti-rabbit lgG H&L (Alexa Fluor®488) secondary antibodies (1:200, Abcam) conjugated with fluorescein were used to visualize the positive stained cells respectively.

### 2.4 Adipogenic differentiation of paNCSCs

A sterile cover slip was inserted into each well of 24-well plate. paNCSCs were collected as described 2.2. Cells were pelleted by centrifuging for 10 minutes with 1000rpm and re-suspended with growth medium at a density of 7.4×10^4^ cells/ml. 0.5 ml of cell suspension were seeded into each well of 24-well plate. After 100% confluent, the growth medium from each well was removed and added adipogenic differentiation medium including fetal bovine serum (10%, Gibco), Pen strep (50 units/mL, Gibco), Amphotericin B (1.25 ug/mL, Gibco), Dexamethasone (200 μmoL/L, Sigma), insulin (5 mg/mL, AbMole), isobutylmethylxanthine (0.4 moL/L, Sigma), indomethacin (1 moL/L, Sigma). Cells were incubated at 37℃ with 5% CO_2_/ 95% air. Change the medium every three days.

### 2.5 Hematoxylin and Oil Red staining

Sterile cover slip of undifferentiation group (C group), 2 days of adipogenic differentiation group (T1 group), 6 days of adipogenic differentiation group (T2 group) were taken, and then stained with hematoxylin (Solarbio) and saturated Oil Red dye (Solarbio). Cells in each group were fixed with formaldehyde calcium for 10 minutes, and washed thoroughly with distilled water twice for 1 minute. Cells were added 60% isopropyl alcohol and immersed for 3 minutes. Cells were stained with saturated Oil Red dye dissolved in distilled water at 3:2 for 10 mintues, and made interstitial transparent with 60% isopropanol (about 30 s-1 min). Cells were counterstained with hematoxylin for 2 mintues and washed with distilled water. The cell morphology was observed light microscopy in 200 randomly selected cells.

### 2.6 RNA-Seq

The cells were passaged into 6-well plates with the method as described in 2.4, cultured with growth medium. Cell samples cleaved by Trizol were were collected. Shanghai OE Biotech Co., Ltd. (Shanghai, China) was commissioned to do RNA Sequencing on days 0, 2, 6 after adipogenesis respectively., The results were performed using the OECloud tools at https://cloud.oebiotech.cn.

### 2.7 Verification of sequencing results

To validate the four adipogenic key genes identified by sequencing, according to the FASTA sequences of C/EBPα (XM_003127015.4), C/EBPβ (NM_001199889.1), C/EBPδ (XM_005663091.2) and PPARγ (NM_214379.1) published in NCBI. Primers were designed separately on Primer 3 Plus and synthesized by Sangon Biotech (Shanghai) Co., Ltd. (Shanghai, China). The primer sequences are shown in Table 1. RT-qPCR was performed according to the Takara SYBR®Premix Ex Taq^TM^ Ⅱ instructions and followed the predenaturation: 95℃ for 30 s, 1 Cycle; PCR reaction was 95℃ for 5 s, 57℃ for 20 s or 58℃ for 20 s, 40 Cycles. Melting curve analysis: 95 ℃ for 0 s, 65 ℃ for 15 s, 95 ℃ for 0 s. Porcine GAPDH gene was used as the internal reference, and the RT-qPCR results were calculated with 2^-ΔΔT^ to calculate the relative expression of the target gene.

**Table 1.**
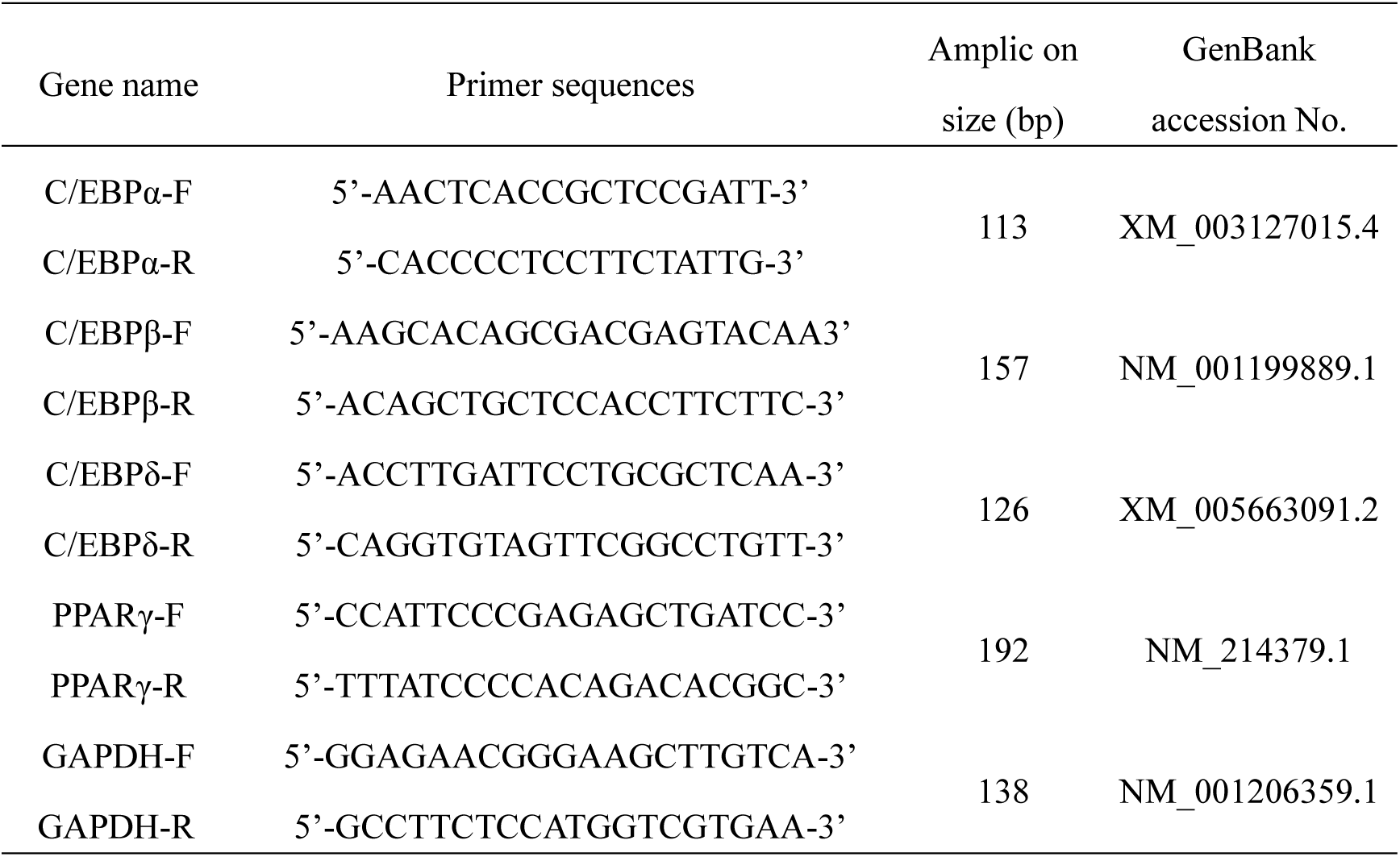
RT-qPCR primer sequences

### 2.7 C/EBPα, δ, β and PPARγ-Specific siRNA (siRNA C/EBPα, δ, β and PPARγ) Transfection

Sangon Biotech (Shanghai) Co., Ltd. (Shanghai, China) was commissioned to synthesize siRNA C/EBPα, δ, β and PPARγ based on pig C/EBPα, δ, β and PPARγ mRNA (Gene Bank XM_003127015.4, XM_005663091.2, NM_001199889.1 and NM_214379.1). RiboFECT^TM^CP Transfection Kit (Guangzhou, China) was used as the transfection reagent. The siRNA sequences are shown in Table 2, and the transfection conditions of siRNA were 20 nM and 50 nM for 0, 2, 4, 6 days.

**Table 2.**
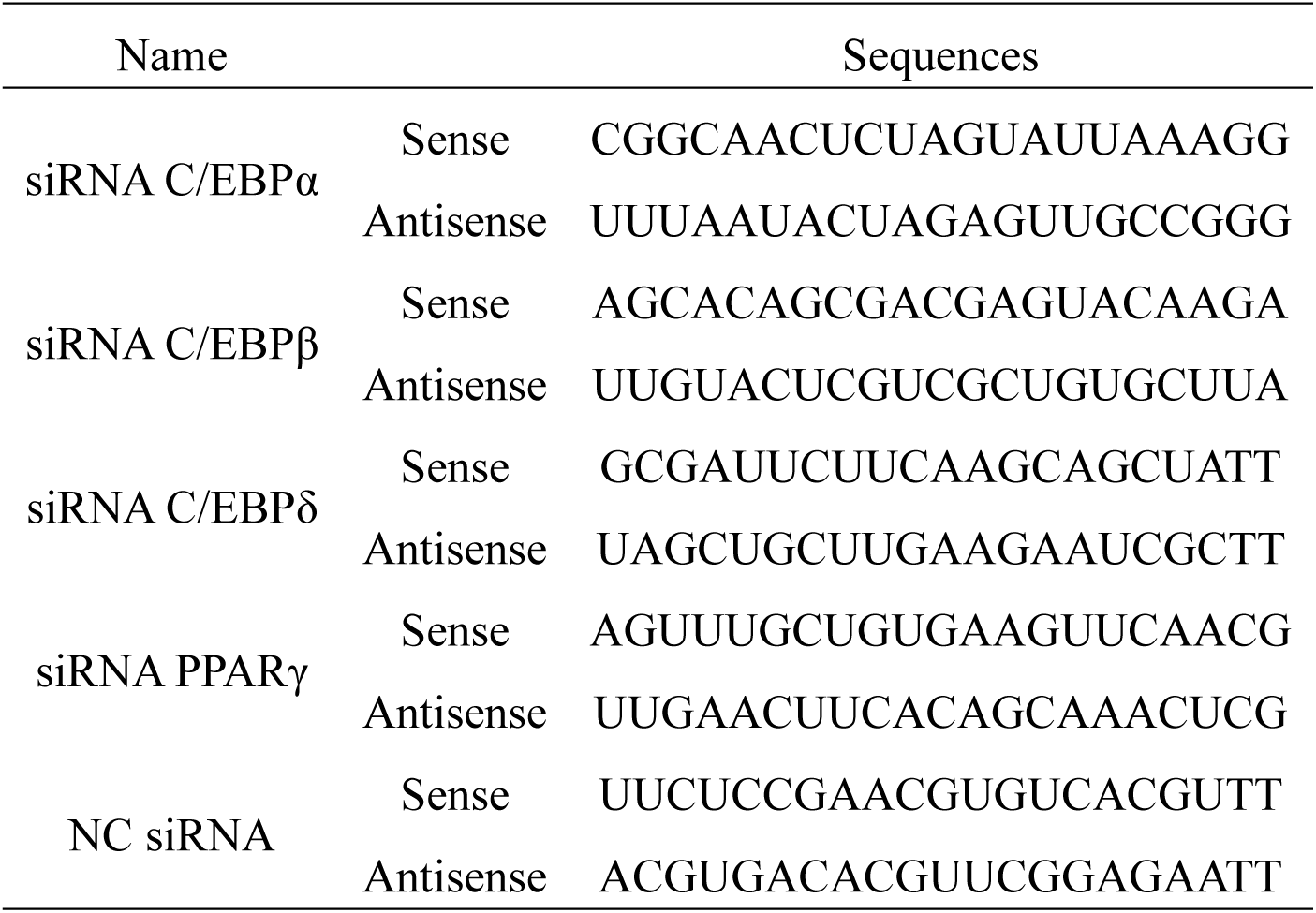
siRNA sequences

### 2.8 RT-qPCR

The culture medium of each group of cells was discarded, and the cells were added with RNAiso Plus (Takara Bio, Japan) to digest. The cells were collected, and total RNA was extracted by Trizol method. RT-qPCR detection was performed on each group of samples by using a reverse transcription kit (Takara Bio, Japan) and fluorescence quantitative kit (Takara Bio, Japan). The primer sequence is the same as Table 1 mentioned in 2.7

### 2.9 Western Blot Analysis

At indicated times of induced and interfered, the culture medium in the cell plate was discarded. The cells were washed three times with PBS and lysed in RIPA buffer containing protease phosphatase inhibitors (Boster biological, Wuhan, China) for 30 min on ice under shaking. The cells were centrifuged at 4℃ and 12000 r/min for 10 min, and the supernatant was collected. Protein concentration was determined using a BCA assay kit (Boster biological, Wuhan, China). The protein sample was mixed with 5 £ protein loading buffer (Boster biological, Wuhan, China) at a ratio of 4:1 and denatured in a boiling water bath for 10 min. The protein samples (25 μg/ lane) were separated using 10% polyacrylamide gels and transferred to 0.45 mm nitrocellulose membranes (PVDF membranes) by using the wet transfer system (Junyi, Beijing, China). After blocking with 5% BSA in TBST for 1 h at 37°C, the PVDF membranes were incubated at 4℃ overnight with primary antibodies for C/EBPα (Abmart, Shanghai, China) and PPARγ (Bioss, Beijing, China). After washing off the primary antibody, the membranes were exposed to goat anti-rabbit IgG-HRP-conjugated antibody (Abmart, Shanghai, China) as secondary antibody for 2 h at 28℃. The secondary antibody was washed, and the ECL chemiluminescence reaction was carried out. Imaging was conducted with a FluorChem HD2 High quality chemiluminescence imaging system (Protein Simple, CA). We used GAPDH (Boster biological, Wuhan, China) as an internal reference protein.

### 2.10 Image and Statistical Analyses

Results are presented as arithmetic mean § standard error. Each experiment was repeated at least three times. The SPSS 22.0 statistical software package (Chicago) was used to perform ANOVA of all data. Histograms were prepared with GraphPad Prism 6.0 software (San Diego, CA). Grayscale value was analyzed by ImageJ software (National Institutes of Health, USA).

## 3 RESULTS

### 3.1 The migration of NCSCs from subcutaneous adipose tissuses

After 2-5 days of explanting, the cells can be seen migrating from fat tissues (**Figure 1A and 1B**), after 9 days of explanting, the cell can be passaged (**Figure 1C**). We examined the expression of p75NTR and SOX2 in the isolated cells. The results show p75NTR (**Figure 1D, 1E, and 1F**) and SOX2 (**Figure 1G, 1H, and 1I**) were expressed in the isolated cells.

**Figure 1.**
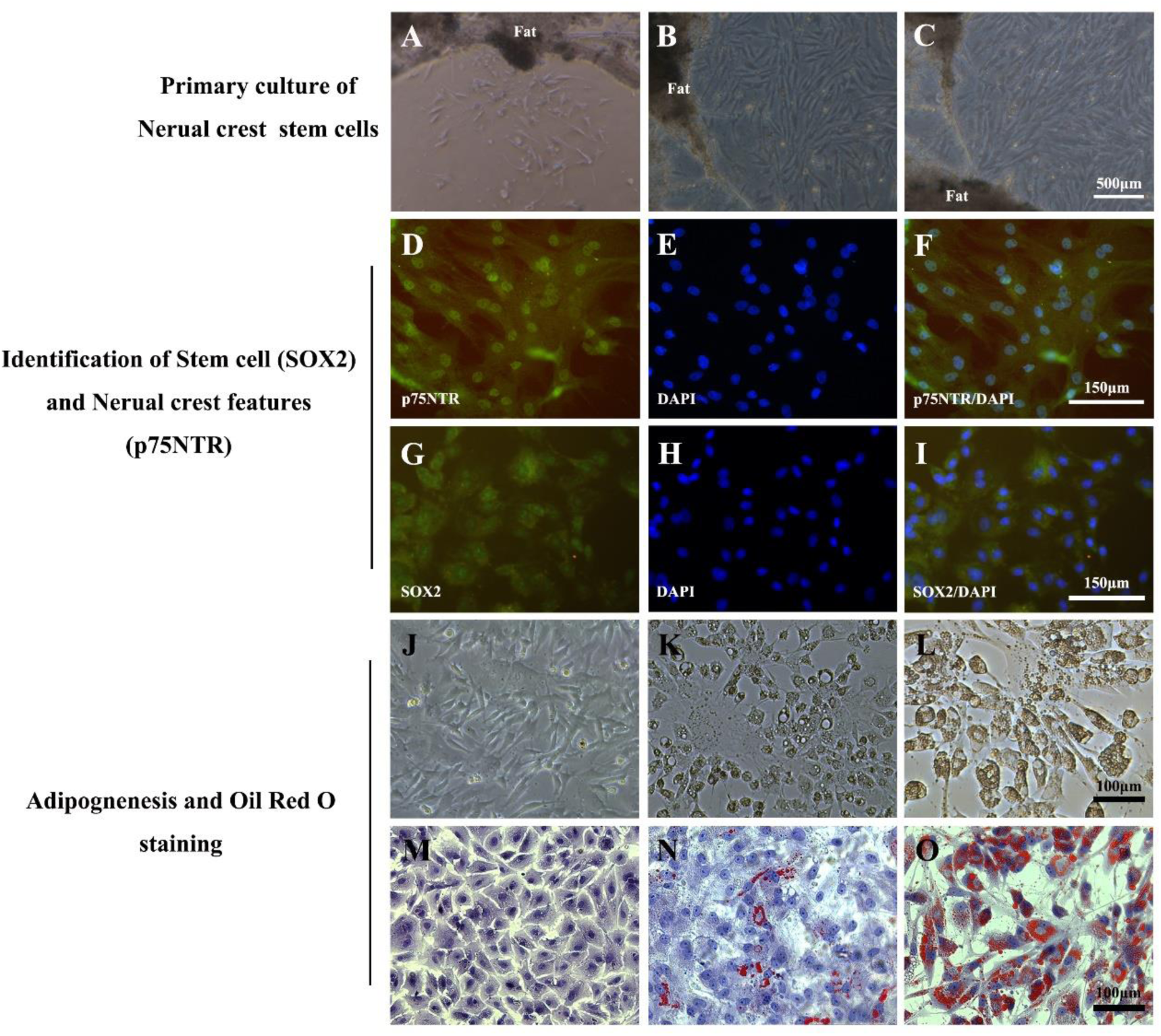
(A, B, and C) The stem cells can be seen migrating from pig fat tissues. (D) The immunofluorescence revealed expression of p75NTR (green) in the isolated cells, (E) nuclear counterstaining was performed using DAPI (blue), (F) the merge of immunofluorescence p75NTR and DAPI. (G) The immunofluorescence revealed expression of SOX2 (green) in the isolated cells, (H) nuclear counterstaining was performed using DAPI (blue), (I) the merge of immunofluorescence SOX2 and DAPI. (J and M) The cells of C group were undifferentiation, (K and N) the cells of T1 group were adipogenic differentiation for 2 days, (L and O) the cells of T2 group were adipogenic differentiation for 6 days.

### 3.2 Identification of adipogenic capability of NCSCs in vitro

The cells of undifferentiation group (C group) were spindle-shaped or polygonal (**Figure 1J**), before the adipogenesis. The cells of 2 days of adipogenic differentiation group (T1 group) were generally vacuolated, and many small dark droplets appeared around the vacuoles in the cytoplasm (**Figure 1K**). the morphology of the cells of 6 days of adipogenic differentiation group (T2 group) were not significantly different from the 2nd day, but the dark droplets in the cytoplasm of the cells were significantly increased (**Figure 1L**). After hematoxylin and Oil Red staining, the small droplets in the cytoplasm of T1 and T2 group were stained red (**Figure 1N and 1O**), that is, fat. Yet, the fat droplets were not seen in the cytoplasm of C group. (**Figure 1M**).

### 3.3 The mRNA sequence analysis of NCSCs before and after adipogenesis

The mRNA sequencing of 9 samples were completed, and obtained 144.23 G clean data. By aligning reads to the reference genome, the genome alignment of each sample was obtained, and the alignment rate was 96.83-98.66%. There were 3 differential gene groups, and the number of differential genes detected were 3755, 4443, 4788. The gene expression profiles of these samples were analyzed with PCA, which revealed significant differences in gene expression patterns from 0 day to 6 days **(Figure.2A)**.

#### 3.3.1 Identifcation of DEGs at different time periods of adipogenesis

We analyzed and compared DEGs between samples before adipogenesis and samples at 2 different stages of adipogenesis. Volcano plots and heat maps were plotted to visualize DEGs and their upregulation and downregulation **(Figures.2B, 2C)**.

To be specific, A total of 1595 genes were up-regulated and 2160 genes were down-regulated between 0 day and 2 days of adipogenesis **(Figure.2B1)**. Compared with 6 days of adipogenesis, 2871 genes were up-regulated and 1572 genes were down-regulated after 2 days of adipogenesis **(Figure. 2B2)**. A total of 2777 genes were up-regulated and 2011 genes were down-regulated between 0 day and 6 days of adipogenesis **(Figure. 2B2)**.

**Figure. 2.**
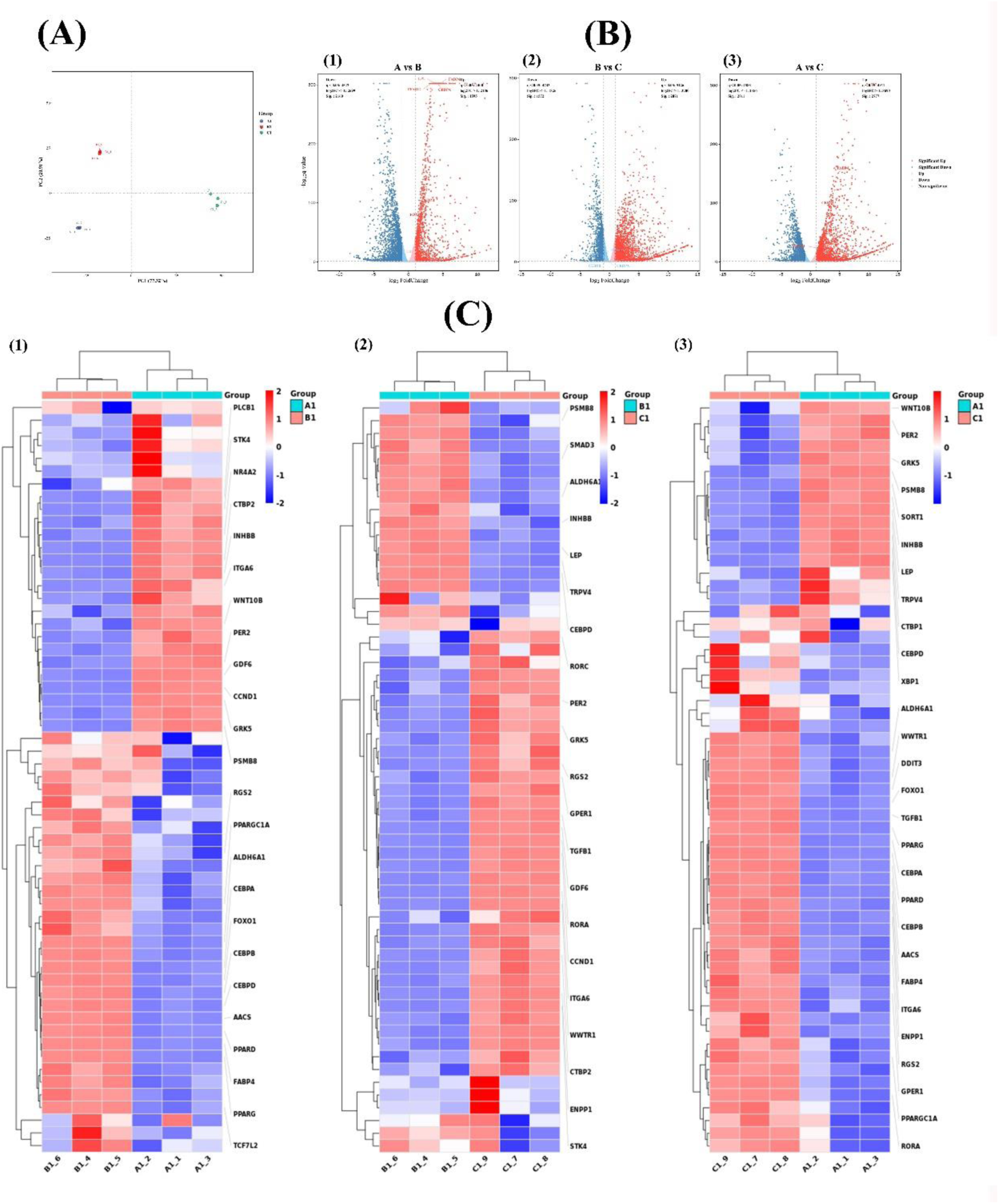
(A) PCA scatter plot of DEGs at 0, 2, 6 days of adipogenesis. Volcano maps of DEGs. Red and blue spots represent DEGs: red spots represent upregulated genes and blue spots represent downregulated genes. B-vs-C, A-vs-C, and A-vs-B represent differentially expressed genes between 2 and 6 days of adipogenesis, 0 and 6 days of adipogenesis, and 0 days and 2 days of adipogenesis, respectively. (C) Heatmap analysis of differentially expressed adipogenic relative genes.

In addition, using adipocyte differentiation as the screening condition, 54, 52 and 29 functional DEGs were screened, respectively. The results were visualized by heatmap. Which showed that 28 genes were up-regulated and 26 genes were down-regulated between 0 day and 2 days of adipogenesis **(Figure. 2C1)**. 20 genes were up-regulated and 32 genes were down-regulated between 2 day and 6 days of adipogenesis **(Figure. 2C2)**. Compared with 0 days of adipogenesis, 18 genes were up-regulated and 11 genes were down-regulated at 6 days of adipogenesis, respectively **(Figure. 2C2)**.

#### 3.3.2 DEGs functional enrichment analysis

The results of GO enrichment of adipogenesis between 0 and 2 days showed that DEGs in the term of molecular function were mainly enriched in nuclear receptor activity and transforming growth factor β receptor binding. For the term of cellular component, DEGs were mainly enriched in the nucleus and transcription repressor complex. For the term of biological process, DEGs were mainly enriched in processes, including white and brown adipocyte differentiation, positive and negative regulation of adipocyte differentiation **(Figure. 3A1)**. In addition, KEGG enrichment analysis showed that DEGs were mainly enriched in Wnt signaling pathway, AMPK signaling pathway and PPAR signaling pathways **(Figure. 3B1)**.

**Figure. 3.**
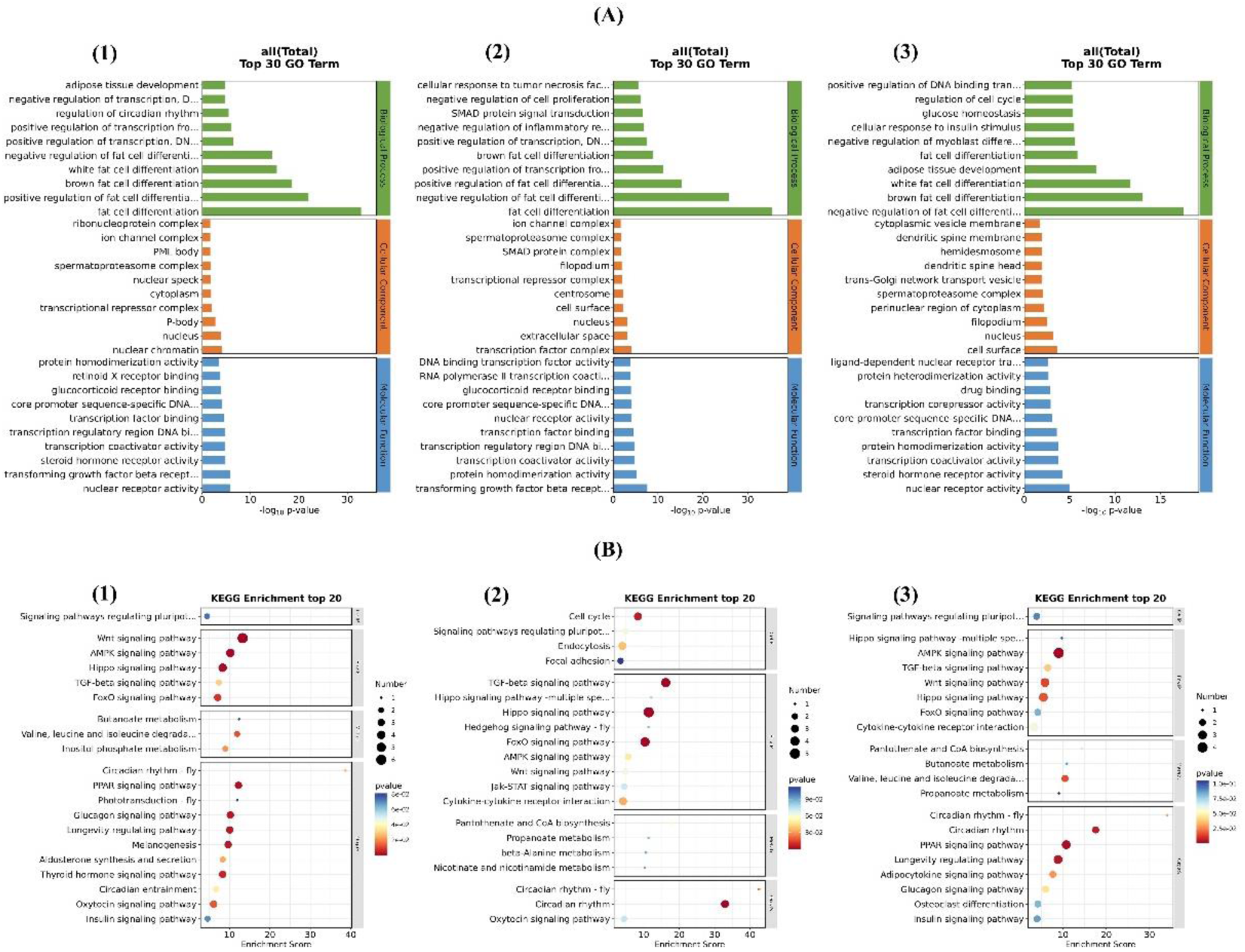
(A) Bar plot of GO enrichment results. (B) Bubble diagram of KEGG enrichment results. Bubble color corresponds to the p value for statistical significance of KEGG pathway enrichment. Bubble size is proportional to the number of genes annotated in a particular pathway.

The results of adipogenesis between 2 and 6 days revealed that the DEGs in the term of molecular function were mainly enriched in protein homodimerization activity and transcriptional coactivator activity. The DEGs in the term of cellular component were mainly enriched in transcription factor complex, extracellular space and nucleus. The DEGs in the term of biological process were similarly enriched in white and brown adipocyte differentiation, positive and negative regulation of adipocyte differentiation, and adipocyte differentiation **(Figure. 3A2)**. In addition, KEGG enrichment analysis showed that DEGs were mainly enriched in TGF-β signaling pathway, Hippo signaling pathway, FOXO signaling pathway, etc **(Figure. 3B2)**.

The results of adipogenesis between 0 and 6 days revealed that the DEGs in the term of molecular function were mainly enriched in RNA polymerase Ⅱ distal enhancer sequence specific binding, kinase binding, kinase activity, etc. The DEGs in the term of cellular component were mainly enriched in RNA polymerase Ⅱ transcription factor complex, heterotrimeric G protein complex, nuclear and plasma. DEGs in the term of biological process were also enriched in white and brown adipocyte differentiation and adipose tissue development **(Figure. 3A3)**. In addition, KEGG enrichment analysis showed that DEGs were widely enriched in a variety of signaling pathways, including AMPK signaling pathway and PPAR signaling pathway **(Figure. 3B3)**

### 3.4 Validation of mRNA expression of adipogenic related genes

Agarose gel electrophoresis of C/EBPα, C/EBPβ, C/EBPδ, and PPARγ common PCR products showed target bands at 113 bp, 157 bp, 126 bp, and 192 bp, respectively, and no non-target bands were amplified (**Figure. 4A**).

**Figure 4.**
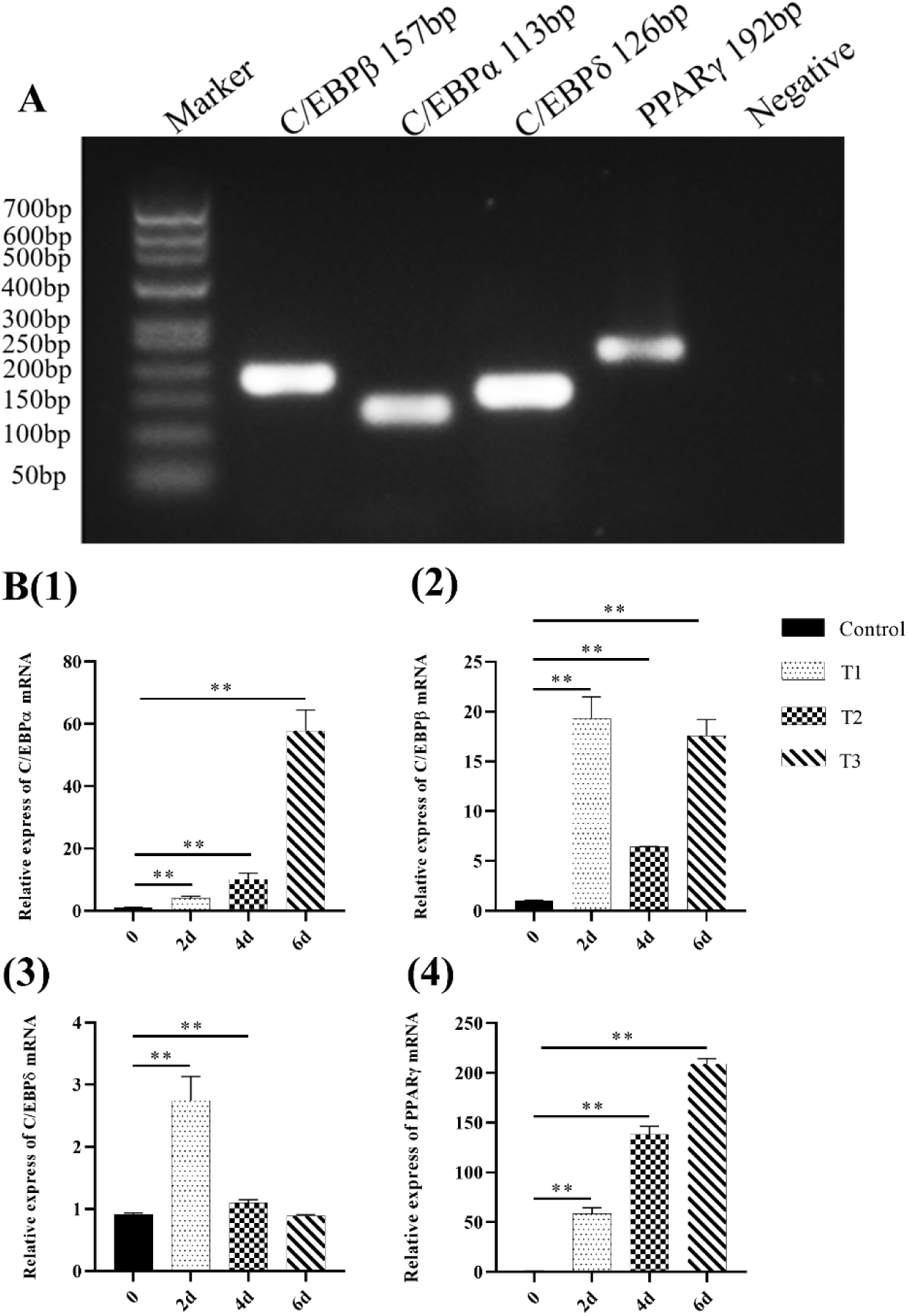
(A) The specific results of PCR. (B1-4) Relative expression of C/EBPα, C/EBPβ, C/EBPδ, and PPARγ mRNA in porcine subcutaneous adipose NCSCs. “**” represents the significant difference (P<0.01).

The relative expression of C/EBPα mRNA in porcine subcutaneous adipose NCSCs was significantly higher than that in control group (P<0.01) after 2, 4, and 6 days of adipogenic induction. The relative expression of C/EBPβ mRNA in group C was significantly higher than that in group C (P<0.01). The relative expression of C/EBPδ mRNA in group C was significantly higher than that in group C (P<0.01). The relative expression of PPARγ mRNA was significantly higher than that in group C (P<0.01) **(Figure. 4B1-4)**.

### 3.5 Regulatory mechanisms of C/EBPα, C/EBPβ, C/EBPδ and PPARγ in adipogenesis

To investigate the expression sequence of C/EBPα, C/EBPβ, C/EBPδ and PPARγ during adipogenesis, specific siRNA targeting the four key genes was used to continuously inhibit their expression from 0 to 6 days of adipogenesis. The relative express levels of C/EBPα, C/EBPβ, C/EBPδ and PPARγ mRNA were detected by RT-qPCR. The relative express of C/EBPα mRNA in T group was significantly higher than that in control group, siRNA C/EBPα group, siRNA C/EBPβ group, siRNA C/EBPδ group and siRNA PPARγ group (P<0.01). There was no significant difference between Negetive control group and T group **(Figure. 5a1-3)**. Similarly, we detected the dynamic changes of C/EBPβ and found that the relative express of C/EBPβ mRNA in T group was significantly higher than that in C group, siRNA C/EBPβ group and siRNA PPARγ group (*P*<0.01), there was no significant difference between siRNA C/EBPα group, siRNA C/EBPδ group, Negetive control group and T group **(Figure. 5 b1-3)**. We detected the dynamic changes of C/EBPδ, and found that the relative express of C/EBPδ mRNA in T group was significantly higher than that in C group, siRNA C/EBPβ group and siRNA C/EBPδ group (*P*<0.01), there was no significant difference between siRNA C/EBPα group, siRNA PPARγ group, Negetive control group and T group **(Figure. 5 c1-3)**. We detected the dynamic changes of PPARγ, found that the relative express of PPARγ mRNA in T group was significantly higher than that in C group, siRNA C/EBPβ group and siRNA PPARγ group (*P*<0.01), there was no significant difference between siRNA C/EBPα group, siRNA C/EBPδ group, Negetive control group and T group **(Figure. 5d1-3)**.

**Figure. 5.**
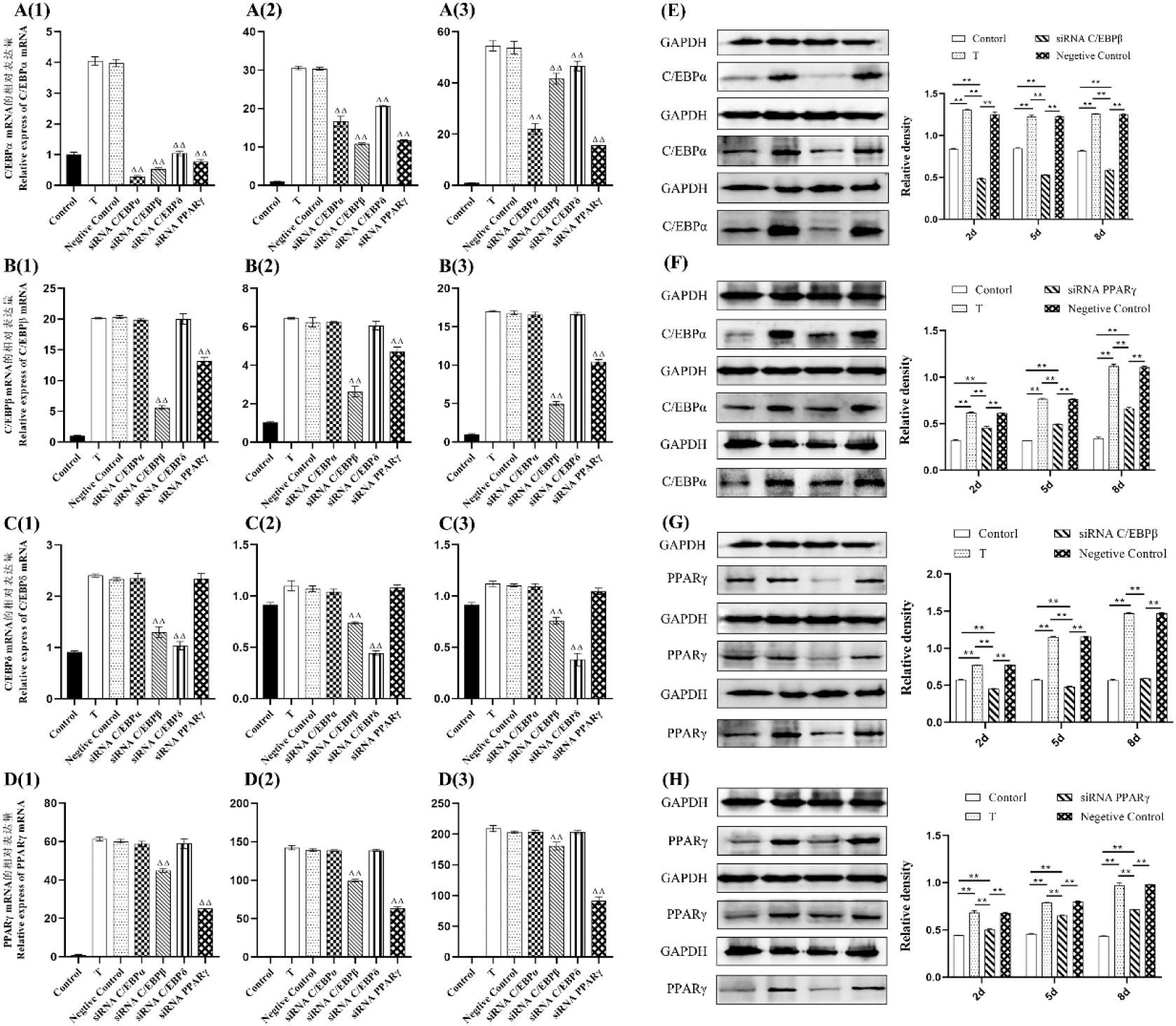
(A1-3) The relative expression of C/EBPα mRNA after 2, 4, 6 days of adipogenesis. (B1-3) The relative expression of C/EBPβ mRNA after 2, 4, 6 days of adipogenesis. (C1-3) The relative expression of C/EBPδ mRNA after 2, 4, 6 days of adipogenesis. (D1-3) The relative expression of PPARγ mRNA after 2, 4, 6 days of adipogenesis. (E) Relative density of C/EBPα protein after 2, 4, and 6 days of adipogenesis and simultaneous transfection of siRNA C/EBPβ. (F) Relative density of C/EBPα protein at days 2, 4, and 6 after adipogenesis and simultaneous transfection of siRNA PPARγ. (G) Relative density of PPARγ protein after 2, 4, and 6 days of adipogenesis and simultaneous transfection of siRNA C/EBPβ. (H) Relative density of PPARγ protein at 2, 4, and 6 day after adipogenesis and simultaneous transfection of siRNA PPARγ. “**” represents the significant difference. “ΔΔ” represents the significant difference.

To verify the RT-qPCR results, paNCSCs were treated with siRNA C/EBPβ and siRNA PPARγ between 0 and 6 of adipogenesis continuously. The results of which relative expression of C/EBPα and PPARγ was detected by western blotting were expressed as relative density **(Figure.5E-H)**. After 2 days of adipogenesis, the protein expression levels of PPARγ and C/EBPα in the T group were significantly higher than those in the Control group and siRNA C/EBPβ group (*P*<0.01). The protein expression levels of PPARγ and C/EBPα in the Control group and the Negative control group were also significantly higher than those in the siRNA groups (*P*<0.01). After 4 days of adipogenesis, the protein expression levels of PPARγ and C/EBPα in the T group were significantly higher than those in the Control group and the siRNA groups (*P*<0.01). The protein expressions of PPARγ and C/EBPα in the Control group and the Negative control group were also significantly higher than those in the siRNA groups (*P*<0.01). After 6 days of adipogenesis, the protein expression levels of PPARγ and C/EBPα in the T group were significantly higher than those in the Control group and the siRNA groups (*P*<0.01). The protein expression levels of PPARγ and C/EBPα in the Control group and the Negative control group were also significantly higher than those in the siRNA groups (*P*<0.01). At the same time, there was no significant difference between T group and Negative control group at three time periods.

## 4. DISUSSION

The ability of migration is found in early neural crest cells [19]. Similarly, high migratory behavior also be showed by adult NCSCs in vitro and in vivo [20]. Mesenchymal stem cells (MSCS) are a type of cell lineage that can continuously grow in vitro and differentiate into adipocytes. [21]. However, no experiment has yet proved the relationship between NCSCs and MSCs. Furthermore, our previous in vitro data prove NCSCs coming from adipose tissue have high phenotypic plasticity. This is also consistent with the previous observation that a large number of trunk NCCs are capable of generating extracellular matrix cell types including adipocytes, neurons, glial cells, as well as pigment cells [8,22]. In our previous study, we successfully isolated and cultured neural crest stem cells (NCSCS) from porcine skin and revealed that NCSCs could differentiate into adipocytes. In this experiment, we focused on the morphological changes of cells. We found that with the change of medium composition (from growth medium to adipogenic induction medium), the cells changed from spindle to round, the intracellular lipid droplets also changed from less to more, and the oil red O staining was red. However, the molecular mechanisms underlying the adipogenic differentiation process of NCSCs remain largely unclear. Subsequently, we will screen DEGs by mRNA sequencing and find out the related adipogenic approches.

As expected, according to PCA results, finding that 3 groups of cell samples were distributed in different regions of two-dimensional space, indicating significant differences in gene function before and after adipogenesis. This difference can also be observed in the volcano plot. Specifically, FABP4, C/EBPα, C/EBPβ, C/EBPδ, PPARγ, FOXO1, LPL were up-regulated between 0 day and 2 days of adipogenesis; after 2-6 days of adipogenesis, C/EBPα, C/EBPβ, PPARγ, LPL, FOXO1, FABP4 genes were down-regulated, and C/EBPδ decreased significantly; FABP4, C/EBPα, C/EBPβ, PPARγ, and LPL genes were all up-regulated, while C/EBPδ was down-regulated between 0 and 6 days of adipogenesis (Figure.4C). Previous studies have shown that C/EBPβ and C/EBPδ are mainly expressed at the early stage of adipocyte differentiation, and their expression levels increase sharply at the early stage of adipocyte differentiation, and their expression levels decrease at the late stage of differentiation, especially C/EBPδ [23]. For C/EBPβ, it may be not completely downregulated, as C/EBPβ can also regulate transcription of C/EBPα and PPARγ at later stages of adipogenesis, which is consistent with the results of this study. Lipidomic researches on the adipogenesis process of 3T3-L1 cells and revealed that FABP4 and C/EBPα expression was increased at the later stage of adipogenic differentiation [24].

In addition, GO enrichment results revealed that DEGs were mainly enriched in the area of biological processes, Including brown and white adipocyte differentiation, positive and negative regulation of fat cell differentiation. The bubble map of KEGG enrichment showed that the functions of DEGs were related to lipid metabolism and adipocyte differentiation. The bubble map of KEGG enrichment showed that the differentially expressed genes were mainly enriched in PPAR signaling pathway, Wnt signaling pathway, FOXO1 signaling pathway, etc. Finally, we determined that the transcription factors C/EBPα, C/EBPβ, C/EBPδ and PPARγ involved in the PPAR pathway were related to adipogenesis, and there might be a regulatory relationship between them.

Kaifan et al [25] found that C/EBPβ and C/EBPδ were highly expressed in the early stage of adipogenesis (48 hours), while PPARγ and C/EBPα were highly expressed in the later stage of adipogenesis (8 days) [26]. Once C/EBPα is expressed, PPARγ and C/EBPα are continuously expressed by binding to their respective C/EBP-regulatory elements, which jointly regulate and induce the adipocyte phenotype [27]. In this study, the RT-qPCR results on 2, 4 and 6 days of adipogenic induction showed that the relative expressions of C/EBPα, C/EBPβ, C/EBPδ and PPARγ mRNA in T group were significantly higher than those in control group (*P*<0.01), indicating that these transcription factors could promote the adipogenesis of NCSCs.

By transfecting specific siRNA of C/EBPβ, C/EBPδ, PPARγ and C/EBPα into cells, we found that C/EBPβ, C/EBPδ and PPARγ regulated C/EBPα transcription, C/EBPβ and PPARγ could regulate each other, and C/EBPβ could regulate C/EBPδ transcription. Interestingly, C/EBPβ induced adipogenesis, and its downregulation blocked adipogenesis [28]. However, Tanaka et al. [29] used gene targeting technology to aquire C/EBPβ(-/-) and δ(-/-) mice. Their embryonic fibroblasts (MEFs) were cultured in vitro and stimulated with lipogenesis, and it was found that C/EBPβ(-/-) and δ(-/-) mice MEFs were impaired lipid formation. In contrast, only C/EBPβ-null had little effect on the adipogenesis of mice. In 3T3-L1 preadipocytes, ectopic expression of C/EBPβ and δ can also promote adipogenesis without giving the cells adipogenic stimulation [30]. Thus, the adipogenic effects of C/EBPβ are different in vitro and in vivo, but only the presence of both C/EBPβ and C/EBPδ can enable cellular adipogenic lineage formation. Except the early roles of C/EBPβ and C/EBPδ, the important role of C/EBPα in lipid differentiation in vitro has been widely confirmed [31], and we found that C/EBPα was regulated by C/EBPβ, C/EBPδ and PPARγ, partial western blot results seem to confirm this. PPARγ has been reported as a precursor adipocyte marker, which is also beneficial to the terminal differentiation of adipocytes [32]. In this study, after 2∼4 days of adipogenic induction and simultaneous transfection with siRNA PPARγ, the relative expression levels of PPARγ mRNA in siRNA PPARγ group were significantly lower than those in T group (*P*<0.01), and found that PPARγ was regulated by C/EBPβ, which was confirmed by western blot. In fact, the C/EBPβ+δ-PPAR-C/EBPα cascade has been demonstrated to promote adipocyte differentiation in many studies, which is in accordance with the results of the present study. Previous research models used 3T3-L1 cells or 3T3-F442A cells, which may lead to differences in expression due to different cell types. However, it is undeniable that C/EBPβ and C/EBPδ play an important role in the early stage of adipogenesis, and PPARγ and C/EBPα play a significant role in the late stage of adipogenesis.

## 5. CONCLUSION

In conclusion, our findings indicate that NCSCs are a type of NCSCs that can differentiate into adipogenic lineage. The DEGs were identified by mRNA sequencing and GO enrichment, and C/EBPα, C/EBPβ, C/EBPδ and PPARγ were found to play an important role in the process of adipocyte differentiation. In addition, NCSCs can also be used as an adipocyte model to investigate the formation mechanism and regulatory mode of obesity and its related metabolic diseases, and provide theoretical basis for its treatment.

